# Probing Notch1-Dll4 Signaling in Regulating Osteogenic Differentiation of Human Mesenchymal Stem Cells using Single Cell Nanobiosensor

**DOI:** 10.1101/2022.04.07.487463

**Authors:** Yuwen Zhao, Rui Yang, Zoe Bousraou, Kiarra Richardson, Shue Wang

**Affiliations:** Department of Chemistry, Chemical and Biomedical Engineering, Tagliatela College of Engineering, University of New Haven, West Haven, CT, 06516, USA; Department of Bioengineering, Lehigh University, Bethlehem, PA, 18015, USA; Department of Biomedical Engineering, University of Connecticut, UConn Health, Farmington, CT, 06030, USA; Department of Biomedical Engineering, Duke University, Durham, NC, 27708, USA

**Author notes:** Corresponding author: Dr. Shue Wang.

**Keywords:** Notch1-Dll4 signaling, osteogenic differentiation, mesenchymal stem cells, microenvironment, 3D spheroids

## Abstract

Human mesenchymal stem cells (hMSCs) have great potential in cell-based therapies for tissue engineering and regenerative medicine due to their self-renewal and multipotent properties. Recent studies indicate that Notch1-Dll4 signaling is an important pathway in regulating osteogenic differentiation of hMSCs. However, the fundamental mechanisms that govern osteogenic differentiation are poorly understood due to a lack of effective tools to detect gene expression at single cell level. Here, we established a double-stranded locked nucleic acid (LNA)/DNA (LNA/DNA) nanobiosensor for gene expression analysis in single hMSC in both 2D and 3D microenvironments. We first characterized this LNA/DNA nanobiosensor and demonstrated the Dll4 mRNA expression dynamics in hMSCs during osteogenic differentiation. By incorporating this nanobiosensor with live hMSCs imaging during osteogenic induction, we performed dynamic tracking of hMSCs differentiation and Dll4 mRNA gene expression profiles of individual hMSC during osteogenic induction. Our results showed the dynamic expression profile of Dll4 during osteogenesis, indicating the heterogeneity of hMSCs during this dynamic process. We further investigated the role of Notch1-Dll4 signaling in regulating hMSCs during osteogenic differentiation. Pharmacological perturbation is applied to disrupt Notch1-Dll4 signaling to investigate the molecular mechanisms that govern osteogenic differentiation. In addition, the effects of Notch1-Dll4 signaling on hMSCs spheroids differentiation were also investigated. Our results provide convincing evidence supporting that Notch1-Dll4 signaling is involved in regulating hMSCs osteogenic differentiation. Specifically, Notch1-Dll4 signaling is active during osteogenic differentiation. Our results also showed that Dll4 is a molecular signature of differentiated hMSCs during osteogenic induction. Notch inhibition mediated osteogenic differentiation with reduced Alkaline Phosphatase (ALP) activity. Lastly, we elucidated the role of Notch1-Dll4 signaling during osteogenic differentiation in a 3D spheroid model. Our results showed that Notch1-Dll4 signaling is required and activated during osteogenic differentiation in hMSCs spheroids. Inhibition of Notch1-Dll4 signaling mediated osteogenic differentiation and enhanced hMSCs proliferation, with increased spheroid sizes. Taken together, the capability of LNA/DNA nanobiosensor to probe gene expression dynamics during osteogenesis, combined with the engineered 2D/3D microenvironment, enables us to study in detail the role of Notch1-Dll4 signaling in regulating osteogenesis in 2D and 3D microenvironment. These findings will provide new insights to improve cell-based therapies and organ repair techniques.

## Introduction

Human mesenchymal stem cells (hMSCs) are adult stem cells isolated from adult tissues such as bone marrow and adipose tissue. hMSCs have great potential in cell-based therapies for tissue engineering and regenerative medicine due to their multipotency and easy accessibility [1, 2]. It has been shown that hMSCs have the potential to differentiate into different lineages, including osteoblasts (bone), neuroblasts (neural tissue), adipoblasts (fat), myoblasts (muscle), and chondroblasts (cartilage) [3, 4]. In particular, osteogenic induction of hMSCs differentiation in conjunction with tissue engineering approaches has been exploited to provide an alternative method besides autologous bone graft to replace or restore damaged bone tissues [5]. This evidence demonstrates the application of hMSCs as a promising cell source in bone tissue engineering, given their regenerative properties. Although the differentiation capacity of hMSCs’ has been studied extensively, the mechanisms that control their plasticity, especially how hMSCs can be differentiated into osteoblasts and make bones, are hindered by the availability of technologies for detecting behaviors of cells with single-cell resolution. Despite conceptual advancements in stem cell differentiation, the cellular and molecular mechanisms that govern hMSCs during osteogenic differentiation remain poorly understood.

Notch signaling is an evolutionarily conserved, intercellular, and contact-dependent signaling pathway which regulates cell fate decisions and plays a significant role during development and organogenesis [5-10]. Many studies have demonstrated the critical role of Notch signaling in cell fate determination, cell proliferation, differentiation, and apoptosis in various cell types [11-14]. Notch signaling pathway is initiated when notch ligand binds to the notch receptor, leading to a cascade of events that eventually cleavage of Notch intracellular domain (NICD). Subsequently, the released NICD then translocates to the nucleus, which regulates transcription of target genes [15]. In mammalian cells, there are multiple ligands that are involved in Notch signaling, for example, Delta-like (Dll1, Dll3, and Dll4) and Jagged (Jag1 and Jag2). Among these five Notch ligands, Dll4 is one of the most notable ligands that are involved in development. In recent years, numerous studies have shown that notch signaling is required for stem cell differentiation, including myogenic differentiation [16], neural stem cell [17], intestinal stem cells [18], hematopoietic stem cells [19], and human embryonic stem cells [20]. For example, previous studies have shown that Notch signaling pathway promote murine mesenchymal stem cells (C3H10T12 cells) differentiation by enhancing BMP9 signaling [21]; Bagheri *et al*. showed that Notch signaling is active during osteogenic differentiation induced by pulsed electromagnetic fields; Wagley *et al*. have showed that canonical Notch signaling is required for BMP-mediated human osteoblast differentiation [22]. However, the fundamental mechanisms of how Notch signaling regulates osteogenic differentiation are unclear due to a lack of effective tools to detect gene expression dynamics at the single cell level. Current approaches for characterizing the regulatory role of Notch signaling in osteogenic differentiation are often limited because they are based on the fixation or physical isolation of cells to identify specific phenotypes. Thus the dynamic behavior of cells is inherently lost when studied in isolation or fixed. Previous studies have shown that Notch signaling has dual effects of induction and inhibition on stem cell osteoblastic differentiation which depends on cell differentiation stage and the time of Notch activation [23]. Therefore, it is essential to monitor the dynamic gene expression in single cells during osteogenic differentiation to elucidate the unrecognized characteristics and regulatory processes of the cells. Moreover, given its importance in stem cell proliferation and differentiation, a better understanding of Notch1-Dll4 signaling in both 2D and 3D microenvironments will provide new insights for stem cell therapy and tissue regeneration.

To address the unmet needs, we present an approach using a double-stranded locked nucleic acid (LNA)/DNA nanobiosensor for monitoring gene expression dynamics during osteogenic differentiation in 2D and 3D microenvironments. In this study, we demonstrated for the first time, the usage of this LNA/DNA nanobiosensor to investigate the role of Notch signaling in regulating hMSCs osteogenic differentiation in 2D and 3D microenvironments. By incorporating this biosensor with live hMSCs imaging during osteogenic induction, we performed dynamic tracking of ALP activity of differentiated hMSCs and Dll4 mRNA expression profiles of during the early stage of osteogenic differentiation. We further investigated the role of Notch1-Dll4 signaling in regulating osteogenic differentiation. Pharmacological perturbation is applied to disrupt Notch1-Dll4 signaling to investigate the molecular mechanisms that govern osteogenic differentiation. Morphological features of osteogenic differentiation, including differentiation efficiency and spheroid sizes, were investigated and compared under different perturbations of Notch signaling. Our results provide convincing evidence supporting that Notch1-Dll4 signaling is involved in regulating hMSCs osteogenic differentiation. For the first time, we identified that Dll4 is a molecular signature of hMSCs osteogenic differentiation. Inhibition of Notch1-Dll4 signaling reduced osteogenic differentiation and enhanced hMSCs proliferation, with increased spheroid size. Taken together, the capability of this LNA/DNA nanobiosensor to probe gene expression dynamics during osteogenesis, combined with the engineered 2D/3D microenvironment, enables us to study in detail the role of Notch1-Dll4 signaling in regulating osteogenesis. Our findings regarding the role of Notch signaling in regulating osteogenesis will provide new insights to improve cell-based therapies and organ repair techniques.

## Results

We established an LNA/DNA nanobiosensor for dynamic gene expression analysis in single hMSC during osteogenic differentiation (**Fig. 1A**). The LNA/DNA nanobiosensor is a double-stranded LNA donor - quencher complex (**Fig. 1A**). The LNA donor probe is a 20-base oligonucleotides sequence with alternating LNA/DNA monomers. The LNA nucleotides are modified DNA nucleotides with higher thermal stability and specificity [24]. The LNA donor sequence is designed to be complementary to the target mRNA sequence. This LNA donor probe will spontaneously bind to the quencher sequence to form a LNA donor-quencher complex. The fluorophore at the 5’ end of the LNA donor is quenched by quencher sequence (Iowa Black Dark Quencher) due to its quenching ability [25]. The LNA donor-quencher complex is then transfected into hMSCs following manufacturers’ instructions. This LNA probe is designed to have a higher binding affinity with the target sequence. Once transfected into cells, in the presence of a target mRNA, the LNA probe is thermodynamically displaced from the quencher, to bind to specific target sequences. This displacement is due to larger difference in binding free energy between LNA probe to target mRNA versus LNA probe to quencher sequence. This displacement reaction permits the fluorophore to fluorescence, thus detecting the gene expression at single cell level (**Fig. 1B-C**).

**Fig. 1.**
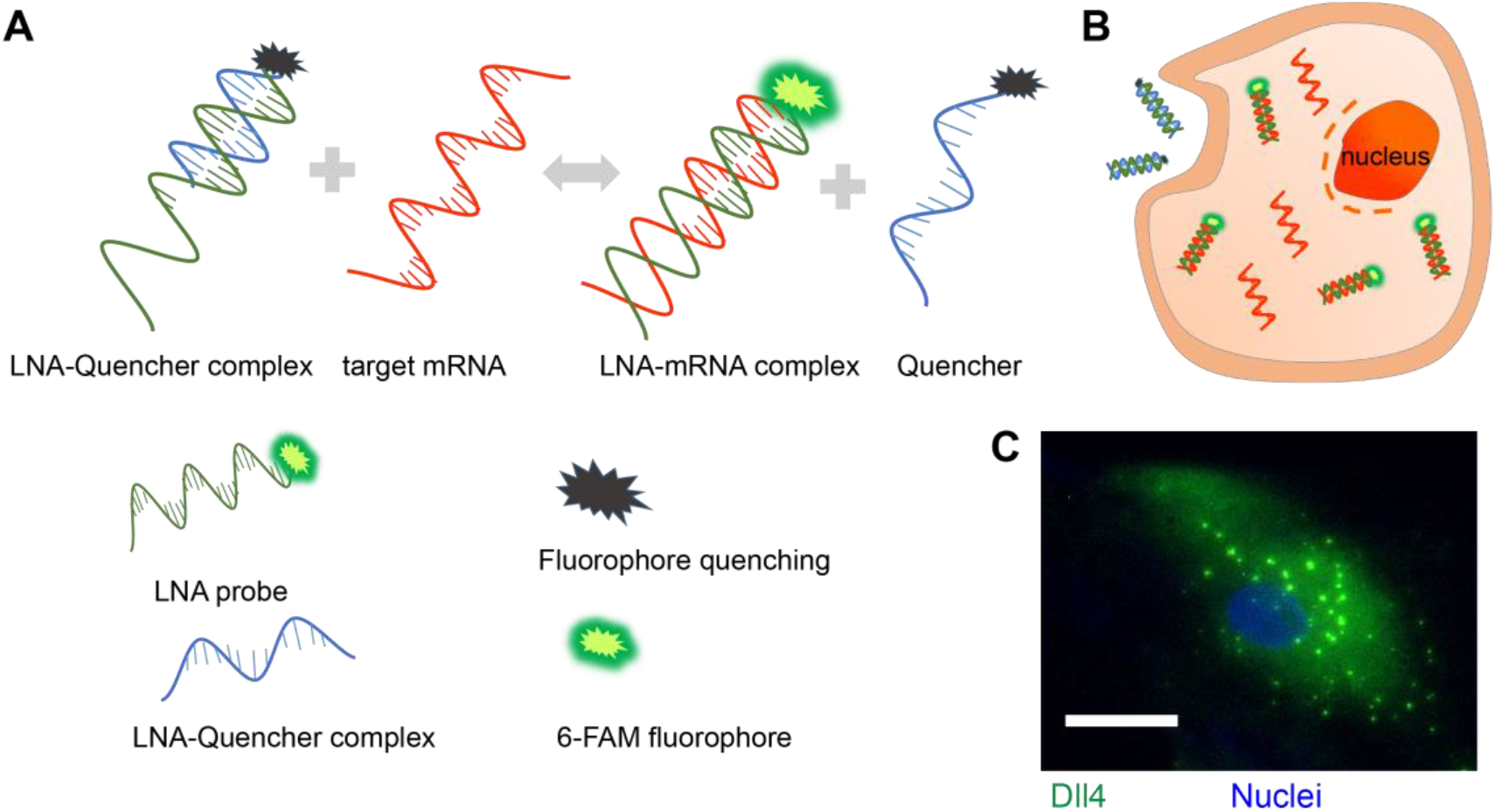
LNA/DNA nanobiosensor for single cell gene expression analysis in living cells. **(A)** Schematic illustration of LNA/DNA nanobiosensor for mRNA detection. Briefly, the LNA/DNA nanobiosensor is a complex of LNA donor and quencher probe. The fluorophore at the 5’ of LNA donor probe is quenched due to close proximity. In the presence of target mRNA sequence, the LNA donor sequence is displaced from the quencher to bind to the target sequence, allowing the fluorophore to fluorescence. **(B)** Schematic illustration of cellular endocytic uptake of LNA/DNA nanobiosensor by cells for intracellular gene detection. **(C)** Representative fluorescence image of Dll4 mRNA expression in single hMSC using LNA/DNA nanobiosensor. Green: Dll4 mRNA; blue: Nuclei. Scale bar: 50 μm.

In order to optimize the LNA/DNA nanobiosensor for monitoring gene expression dynamics during osteogenic differentiation in both 2D and 3D microenvironments, we first characterized the optimal quencher-to-donor ratio to minimize the background noise caused by free fluorophore during the reaction. The donor concentration was set to 100 nM. The quencher concentrations were adjusted such that the quencher-to-donor ratio ranges from 0.5 to 10. The fluorescence intensity was then measured at different quencher-to-donor ratios. As shown in **Fig. 2A**, the fluorescence intensity decreases as the quencher-to-donor ratio increases. To acquire the optimal quencher-to-donor ratio to minimize background noise, we further calculated the quenching efficiency at different ratios. Quenching efficiency was determined by subtracting the fluorescence intensity of free fluorophore by the fluorescence intensity of the donor-quencher complex, divided by the fluorescence intensity of free fluorophore, multiplying the result by 100. Thus, quenching efficiency were calculated as 28.7%, 57.7%, 87.8%, 97.1%, 97.6%, 98.1%, 98.3%, and 98.6%, as the quencher-to-donor ratios were 0.5, 1, 1.5, 2, 4, 6, 8, and 10, respectively. This result indicates that the quenching efficiency increases as quencher-to-donor ratio increases. The intensity of fluorophore was quenched to a very low level, about 3% of the maximum intensity level, at a quencher-to-donor ratio of 2. Further increase of the quencher concentrations did not significantly reduce the fluorescence intensity, consisting to previously reported results [26, 27]. Thus, we set the quencher-to-donor ratio to 2 for the subsequent studies. We next characterized the detectable range of target concentrations of this LNA/DNA nanobiosensor. The fluorescence intensity was measured by varying the DNA target oligonucleotide concentrations while setting the LNA probe concentration at 100 nM. As shown in **Fig. 2B**, the sigmoid-shaped titration curve shows a large dynamic range for quantifying target concentrations ranging from 1 nM to 1000 nM, indicating this LNA probe provides a sufficient large dynamic range for detecting target mRNAs. We further examined the background fluorescence intensity of this LNA probe with different culture mediums and did not observe non-specific binding in different conditions including hMSCs culture and induction medium, **Fig. 2C**. Moreover, we evaluated the stability of this LNA/DNA nanobiosensor by incubating this probe with target mRNA for different duration, ranging from 1 to 14 days. As shown in **Fig. 2D**, the fluorescence intensity did not show a significant difference as the incubation time increased, indicating that the stability of this LNA probe is not affected by incubation time. All these results suggest that the fluorescence of this LNA probe is specific, sensitive, and stable for target mRNA detection.

**Fig. 2.**
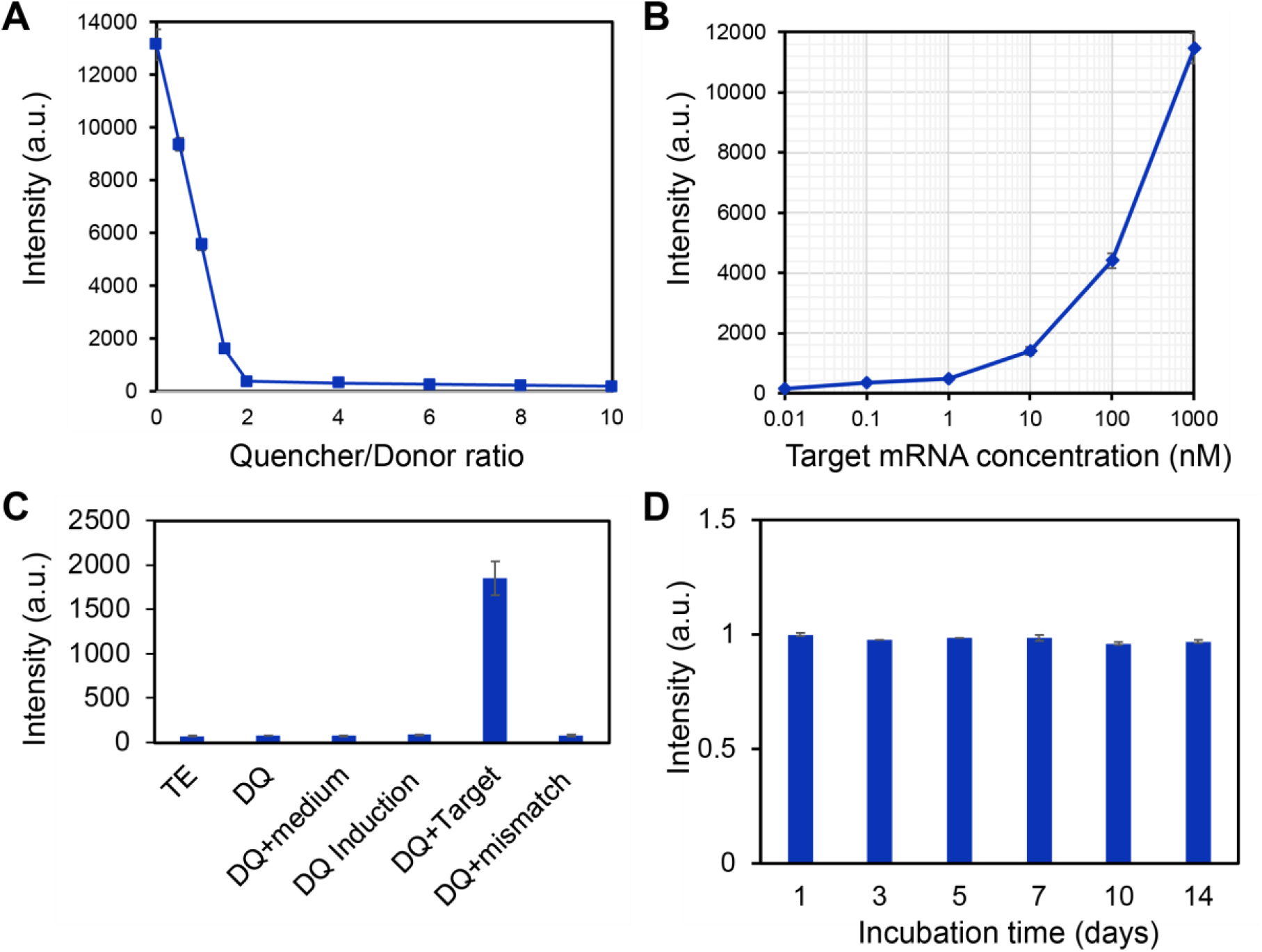
Characterization of LNA/DNA nanobiosensor. **(A)** Calibration of the quencher-to-donor ratio. Fluorescence intensity were measured using a microplate reader by adjusting the quencher concentration. The donor concentration was set to 100 nM. **(B)** Calibration of the detection dynamic range of the LNA/DNA nanobiosensor by varying the target sequence concentration. The concentration of LNA/DNA probe was set to 100 nM. **(C)** Fluorescence intensity of LNA/DNA nanobiosensor in different buffer. TE: Tris-EDTA buffer; medium: hMSCs basal culture medium; induction: osteogenic induction medium; target: target sequence strand were added; mismatch: mismatched target sequence. **(D)** Stability of LNA/DNA nanobiosensor over the course of 14 days. Data are expressed as mean ± SEM. Experiments were repeated three times independently with triplets.

Osteogenic differentiation is a dynamic cellular process that consists of distinct sub-stages and dynamic transcriptional responses during hMSCs differentiation into osteogenic or adipogenic lineages [28]. It has been reported that there are three distinct early stages of cell fate determination during osteogenic differentiation. These three stages can be identified and determined by transcription factor dynamics when cells adopt a committed phenotype: initiate differentiation (0-3 hr, phase I), acquire lineage (4-24 hr, phase II), and early lineage progression (48-96 hr, phase III). Although previous gene expression studies have identified several critical regulatory signaling pathways that are involved in hMSCs early osteogenic differentiation, it is obscure how the gene expression dynamics relate to early osteogenic different progression. To examine the early osteogenic differentiation process, we monitored the transcription factor Dll4 mRNA expression dynamics using the LNA/DNA nanobiosensor. A random probe was included as a control, **Tab. S1**. Both random and Dll4 LNA probes were transfected one day before osteogenic induction. The gene expression profile was determined by measuring fluorescence intensity for each cell daily for 7 days and calculated using NIH ImageJ software. We first confirmed random probe has minimum fluorescence background in hMSCs that were cultured in both basil and osteogenic differentiation medium, **Fig. 3A, Fig. S1**. There is no significant difference in fluorescence intensity of hMSCs between the control group and osteogenic differentiated group, and among each group, **Fig. 3C**. Compared to the random probe, an increase in Dll4 mRNA expression was observed for all the cells in the osteogenic induction group. **Fig. 3B** and **Fig. S2** showed the representative bright and fluorescence images of hMSCs after 3, 5, and 7 days of osteogenic induction. The yellow arrows indicate expressed Dll4 mRNA in osteogenic differentiated hMSCs. We first observed that the Dll4 mRNA expression level was increased in osteogenic induced hMSCs compared to hMSCs cultured in basil medium. This finding is consistent with previous results reported by other groups [29]. To further compare the Dll4 mRNA expression levels, we quantified Dll4 mRNA expression dynamics by measuring the mean fluorescence intensity of each hMSC. Interestingly, after osteogenic induction, the Dll4 mRNA expression was increased and reached a peak after 5 days of induction, then the Dll4 expression was observed to decrease and stayed at a stable level after 7 days of induction, **Fig. 3D**. Dll4 mRNA expression level was increased 4 folds from day 1 to day 5. After 5 days of osteogenic induction, the Dll4 mRNA expression level was reduced one-fold and kept at the same level for 6 and 7 days of induction. These results indicate that Dll4 mRNA expression dynamics indicate two different stages during osteogenic differentiation: early differentiation stage (day 1-day 5) and calcium mineralization stage (after day 5). It has been reported that the process of hMSCs osteogenic differentiation includes early lineage progression (proliferation), early differentiation, and later stage differentiation (calcium mineralization) [28, 30-32]. Our results indicate that Dll4 mRNA expression could potentially be the molecular signature of early osteogenic differentiation.

**Fig. 3.**
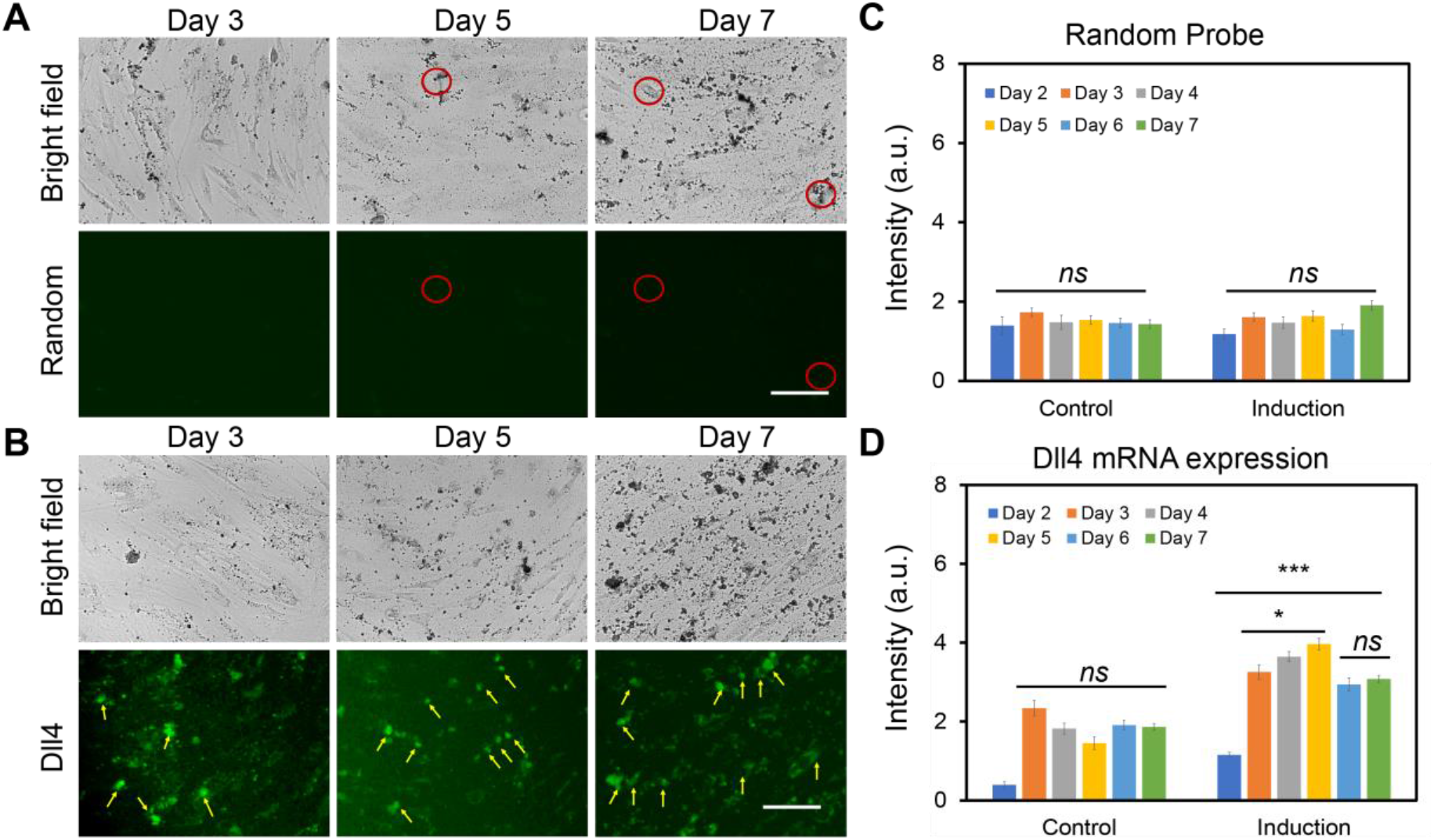
Negative control and Dll4 mRNA tracking of hMSCs during osteogenesis. Representative bright field and fluorescence images of hMSCs after 3 d, 5 d, and 7 d of osteogenic induction. **(A)** Green fluorescence indicates the signal of negative control of random probe. **(B)** Dll4 mRNA expression tracking of hMSCs during osteogenic induction after 3 d, 5 d, and 7 d of osteogenic induction. Green fluorescence indicates Dll4 mRNA expression. Yellow arrows indicate the differentiated cells expressing Dll4 mRNA. **(C)** Mean fluorescent intensity of random probe in hMSCs at different time points under control and induction group. **(D)** Mean fluorescent intensity of Dll4 mRNA expression in hMSCs at different time points under control and induction group. Control group: cells were cultured and maintained in basal medium. Induction group: Cells were cultured in basal medium and induced by osteogenic induction medium after 2 days of cell seeding. Images are representative of at least three independent experiments. Scale bar: 50 μm. Data represent over 100 cells in each group and are expressed as mean ± s.e.m. (n = 3). *p*-Values were calculated using a two-sample *t*-test with respect to day 2. *, *p* < 0.05; **, *p* < 0.01; ***, *p* < 0.005.

The above results showed that Dll4 mRNA expression might relate to hMSCs osteogenic differentiation. Thus, we hypothesize that Dll4 mRNA is a molecular signature of osteogenic differentiated hMSCs. To test our hypothesis, we simultaneously detected Dll4 mRNA expression and alkaline phosphatase (ALP, a biochemical marker for bone formation) enzyme activity during hMSCs osteogenic differentiation using LNA/DNA nanobiosensor and ALP staining assay (Sigma Aldrich), **Fig. 4**. We performed the dsLNA probe assay and immunostaining to correlate differentiated cell characteristics with the expression of Dll4 mRNA. A random probe was utilized as a negative control. The Dll4 mRNA expression and ALP enzyme activities were imaged and quantified after 5 and 10 days of osteogenic induction, respectively. **Fig. 4A** showed representative images of hMSCs after 5 days of culture in basal medium (upper panel) and osteogenic induction medium (lower panel), respectively. Compared to undifferentiated cells, differentiated cells showed enhanced ALP activity and high expression of Dll4 mRNA. In contrast, undifferentiated cells in the control group and induction group showed low levels of Dll4 mRNA expression and ALP activities. Differentiated cells showed high levels of ALP activity, while with the random probe, there is no significant signal detected in hMSCs, **Fig. S3**. We next analyzed ALP activity in hMSCs in both control and induction groups. **Fig. 4B** showed the comparison of ALP activity after 5 days and 10 days of culture, respectively. Compared to hMSCs cultured with basal medium, ALP activity in hMSCs cultured in osteogenic induction medium increased 3.2 folds and 6.5 folds after 5 and 10 days of induction, respectively. We further measured and quantified Dll4 mRNA expression of hMSCs after 5 days and 7 days of osteogenic induction. Dll4 mRNA expression levels were increased by 5.2 folds and 4.56 folds after 5 days and 10 days of osteogenic induction, compared to the control group, **Fig. 4C**. These results showed that Dll4 mRNA is correlated to ALP activity, which is a molecular biomarker of differentiated cells. Moreover, Dll4 mRNA expression showed a dynamic profile during osteogenic differentiation. Meanwhile, the intensity of the random probe maintained a low level regardless of differentiated or undifferentiated cells. Taken together, these results indicate that Dll4 is a molecular signature of osteogenic differentiation of hMSCs.

**Fig. 4.**
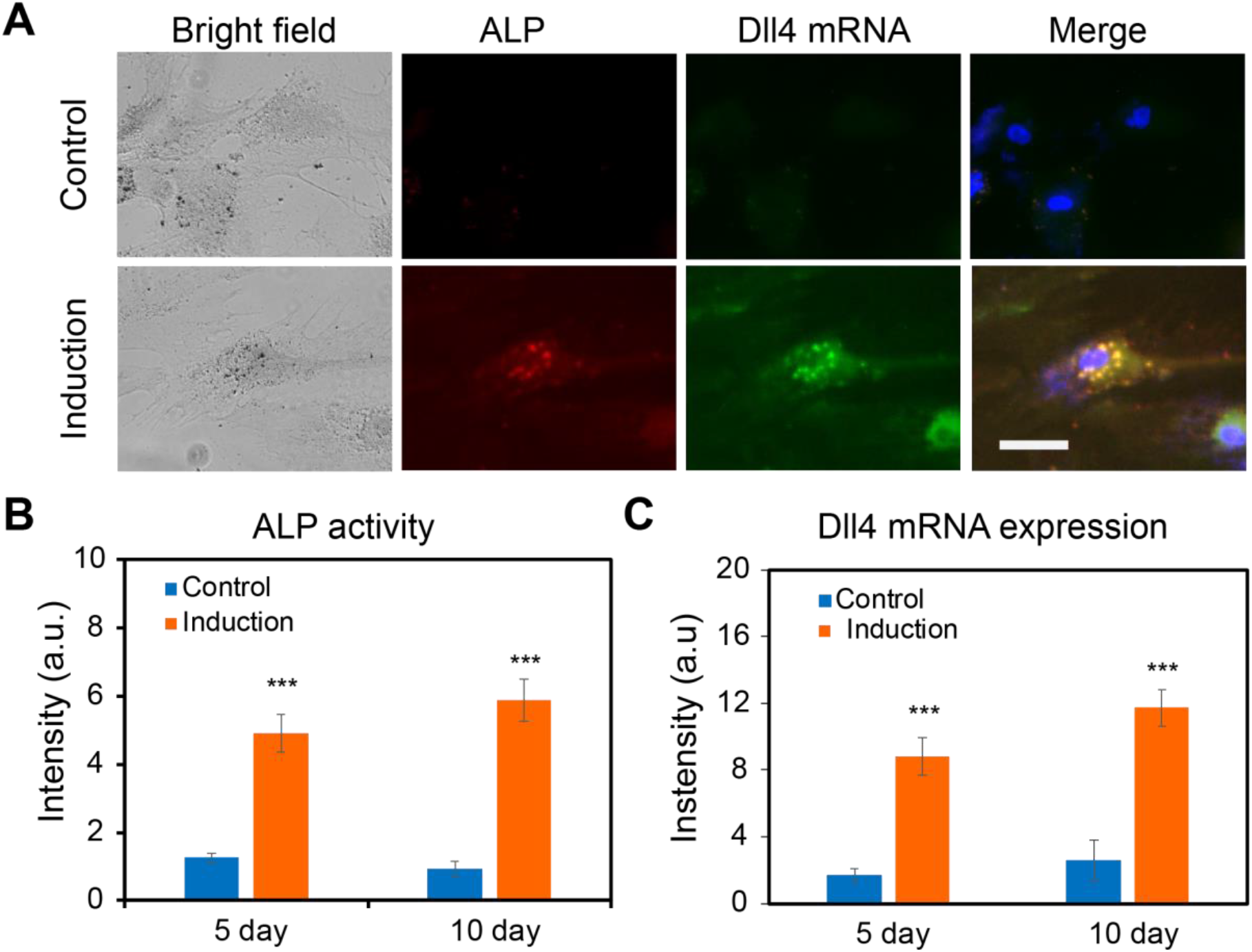
Dll4 mRNA is a molecular signature of hMSCs differentiation. **(A)** Representative bright filed and fluorescent images of single hMSCs under control and induction groups after 5 days. Green: Dll4 mRNA; red: ALP activity; blue: nucleus. Control group: cells were cultured and maintained in the basal medium. Induction group: Cells were cultured in basal medium and induced by osteogenic induction medium after 2 days of cell seeding. Images are representative of at least three independent experiments. **(B-C)** Mean fluorescent intensity of ALP activity and Dll4 mRNA expression after 5 days and 10 days of hMSCs osteogenic induction. ALP activity **(B)** and Dll4 expression **(C)** comparison in the control and induction groups, respectively. Data represent over 100 cells in each group and are expressed as mean± s.e.m. (n=4, ***, *p*<0.001, **, *p*<0.01) Scale bar: 100 μm.

In order to investigate the involvement of Notch1-Dll4 signaling during hMSCs osteogenic differentiation, we perturbed Notch1-Dll4 signaling using two pharmacological drugs, DAPT and Jag1 peptide. DAPT is a γ-secretase inhibitor that blocks Notch endoproteolysis and thus serves as a Notch signaling inhibitor [33]. Jag1 peptide can activate Notch signaling by inhibiting the function of endogenous Jag1, a Notch ligand that has weak signaling capacity but competes with Dll4 [34]. hMSCs were treated with DAPT (20 μM) and Jag1 (40 μM) during osteogenic differentiation to examine the drug effects. A control group was designed without osteogenic induction. The osteogenic differentiation and Dll4 mRNA expression under different treatments were evaluated and compared. Osteogenic differentiation after induction was evaluated by quantifying ALP activity under different treatments. Our results showed pharmacological treatments with DAPT and Jag1 effectively induced changes in osteogenic differentiation, **Fig. 5A**. Specifically, DAPT treatment inhibited osteogenic differentiation and decreased Dll4 mRNA expression, **Fig. 5A**. After 5 days of osteogenic induction, ALP activity in the DAPT treatment group was mediated compared to the induction group without treatment. Particularly, the ALP enzyme activity decreased by 29.9% ((ALP fluorescence intensity in induction group – ALP fluorescence intensity in the presence of DAPT) / ALP fluorescence intensity in induction group) compared to the osteogenic induction group, **Fig. 5C**. These results indicate that inhibition of Notch1-Dll4 signaling using γ-secretase inhibitor mediates osteogenic differentiation of hMSCs. Interestingly, with the treatment of Jag1, osteogenic differentiation efficiency did not show a significant difference, **Fig. 5C**. However, it has been reported that Notch1 ligand, Jag1 has been demonstrated to be essential in various development processes, including osteogenesis. Deng *et al*. recently reported that immobilized Jag1 mimetic ligand activates Notch signaling via the upregulation of NICD, leading to the enhanced stem cell osteogenesis.[35] Jag1 mediated Notch signaling activation is essential for stem cell proliferation and differentiation [36, 37]. A previous study showed that exposure of mouse MSC to Jag1 inhibited osteogenesis, indicating the complex role of Jag1 during stem cell differentiation [37].

**Fig. 5.**
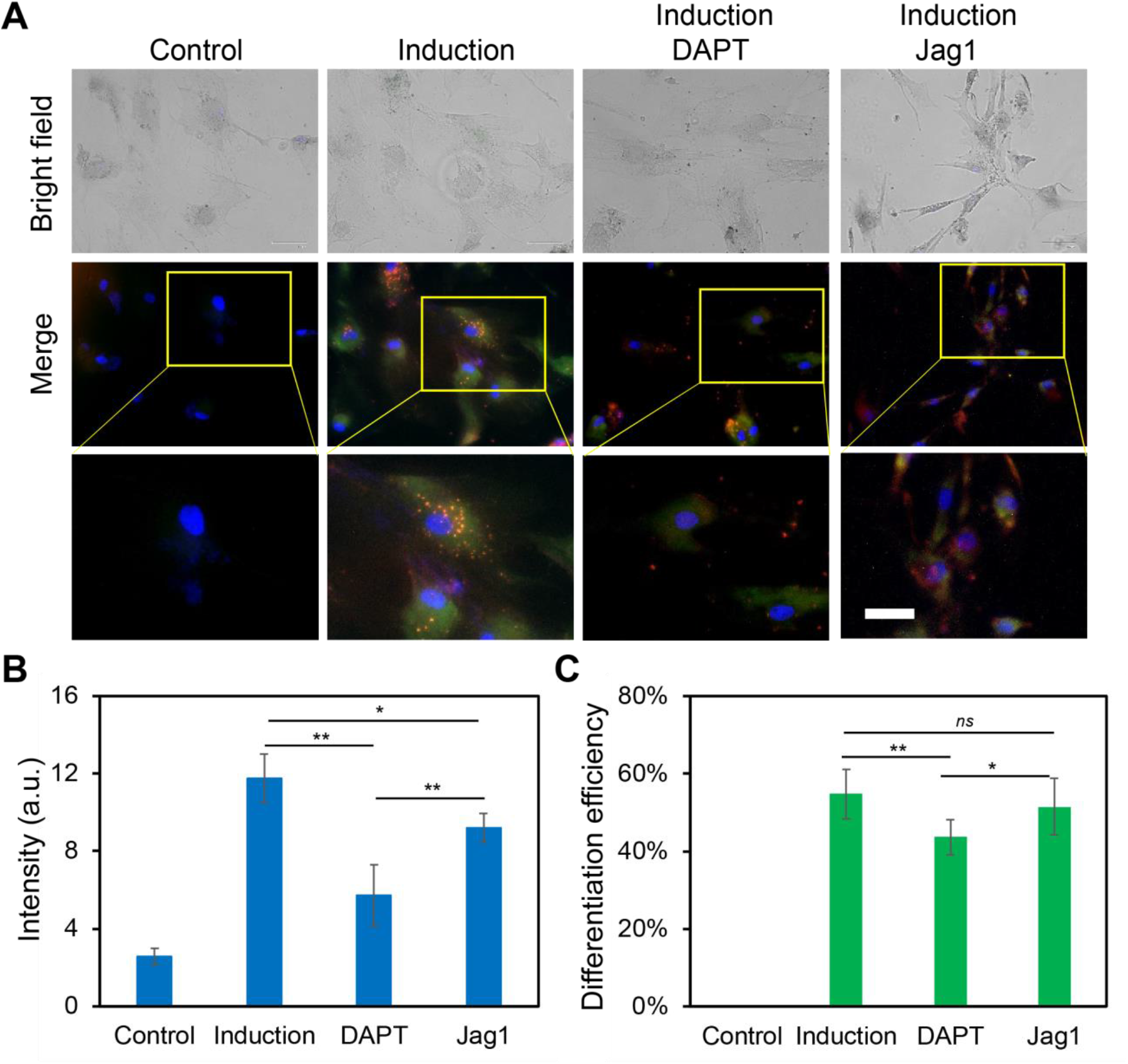
Notch1-Dll4 signaling regulates hMSCs osteogenic differentiation. **(A)** Representative bright field and fluorescence images of hMSCs in control, induction, DAPT and Jag1 treatment groups. Control group: cells were cultured and maintained in basal medium. Induction group: Cells were cultured in basal medium and induced by osteogenic induction medium after 2 days of cell seeding. DAPT group: cells were treated with DAPT (20 μM) daily. Jag1 group: cells were treated with Jag1 (40μM) daily. The bottom images are enlarged areas of a single hMSC. Green: Dll4 mRNA expression; red: ALP activity; blue: nucleus. Scale bar: 100 μm. **(B)** Mean fluorescence intensity of Dll4 mRNA expression under different treatments. **(C)** Comparison of hMSCs osteogenic differentiation efficiency under different treatments. Data represent over 100 cells in each group and are expressed as mean± s.e.m. (n=4, ***, *p*<0.001, **, *p*<0.01)

To further investigate the effects of Notch1-Dll4 signaling during hMSCs induced osteogenic differentiation, we examined Notch1 ligand, Dll4 mRNA expression under different treatments using LNA/DNA probe. Compared to the control group, hMSCs cultured with the osteogenic induction medium showed a significant increase (∼ 4 folds) in the expression of Dll4 mRNA. Meanwhile, with the treatment of γ- secretase inhibitor DAPT, a significant decrease of Dll4 mRNA (32.7%) was observed compared to the osteogenic induction group, **Fig. 5B**. Jag1 treatment also reduced the Dll4 mRNA expression compared to the induction group. However, compared to the DAPT group, Dll4 expression was increased with the treatment of Jag1. These results suggest a positive correlation between Dll4 mRNA expression and osteogenic differentiation. As the Notch pathway is triggered by binding specific ligands to receptors, our results indicate that the expression of Notch ligand Dll4 was modulated during osteogenic differentiation. Dll4 mRNA was significantly increased at the end of the differentiation period (**Fig. 5B**). Notch inhibitors induced a significant decrease of ALP enzyme activity and Dll4 mRNA expression compared to osteogenic induction.

Recently, there has been a growing interest in understanding hMSCs differentiation in 3D microenvironments. It has been shown that the self-assembly of hMSCs into tightly packed clusters of cells in each aggregate mimic “*in vivo*-like” microenvironment and preserves hMSCs phenotype and innate properties [38, 39]. Moreover, it has been reported that the formation of 3D aggregates, or spheroids enhanced the regenerative capacity of hMSCs by promoting the secretion of proangiogenic and chemotoxic factors, and improved cell retention, and survival in preclinical studies [40]. Here, we investigated the effects of Notch1-Dll4 signaling in regulating osteogenic differentiation of 3D hMSCs spheroids. We first fabricated 3D self-organized spheroids of hMSCs using the hanging drop approach [41]. The self-organized 3D spheroids were collected and seeded in Matrigel, **Fig. S5**. To reveal the mechanisms of Notch1-Dll4 signaling, 3D hMSCs spheroids were treated using DAPT and Jag1, respectively. A control group without osteogenic induction was designed for comparison. **Fig. 6A** showed the representative images of hMSCs spheroids under different treatments. Compared to the control group, hMSCs spheroids cultured in osteogenic induction medium showed increased ALP activity and Dll4 mRNA expression, **Fig. 6A-B**. Both DAPT and Jag1 treatments inhibited osteogenic differentiation of 3D hMSCs spheroids. We further quantified the ALP activities and the size of spheroids under different treatments. Compared to the control group, ALP activity was increased ∼ 18 folds for the spheroids in the osteogenic induction group. With Notch pathway inhibitor DAPT treatment, the expression of ALP was significantly decreased, with a 54.2% decrease compared to the induction group. Meanwhile, Jag1 treatment resulted in a 30.2% decrease compared to the induction group. We next examined the sizes of the spheroids under different treatments. Without osteogenic induction, the diameter of the 3D spheroids is approximately 266.5 μm, **Fig. 6C**. The diameter of the spheroids was decreased by 18.6% in the osteogenic induction group. In contrast, compared to the induction group, the diameter of the spheroids was increased by 33.3% and 31.3% with the treatments of DAPT and Jag1, respectively, **Fig. 6C**. These results indicated that inhibition of the Notch1-Dll4 pathway by γ-secretase inhibitor DAPT could inhibit osteogenic differentiation and enhance the proliferation of hMSCs spheroids.

**Fig. 6.**
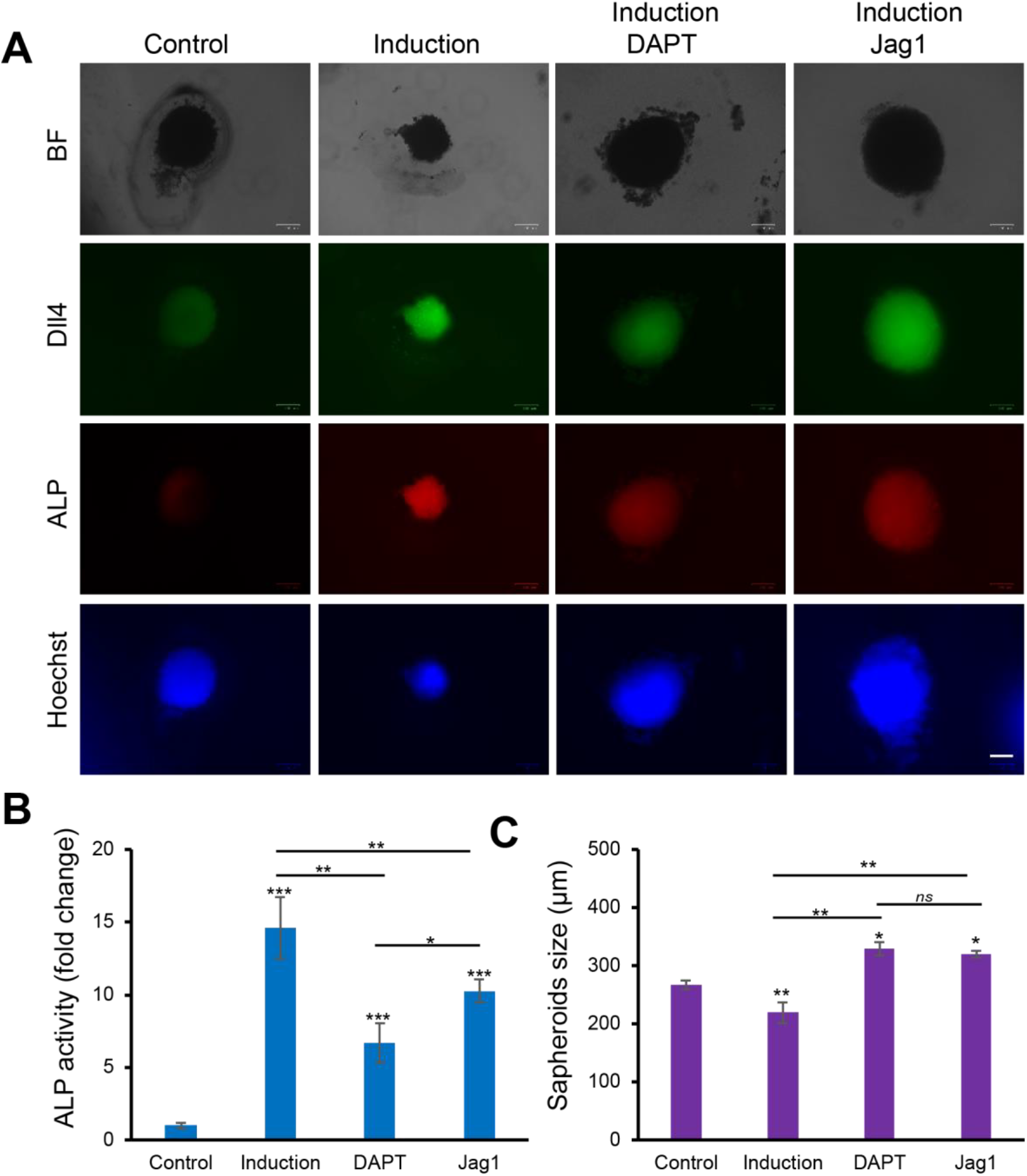
Notch1-Dll4 signaling regulates 3D hMSCs spheroids osteogenic differentiation. **(A)** Representative bright field and fluorescence images of hMSCs spheroids in control, induction, DAPT and Jag1 treatment groups. Green: Dll4 mRNA expression; red: ALP activity; blue: nucleus. Scale bar: 100 μm. **(B)** Quantification of ALP activity of hMSCs spheroids under different treatments. **(C)** Characterization of spheroids sizes under different treatments. Data represent over 50 spheroids in each group and are expressed as mean± s.e.m. (n=4, ***, *p*<0.001, **, *p*<0.01)

## Discussion

In this study, the LNA/DNA nanobiosensor is exploited to monitor the mRNA expression of hMSCs during osteogenic induction. Unlike other conventional reverse transcription polymerase chain reaction (RT-PCR) techniques, this method only requires a small number of cells without lysis. The LNA/DNA nanobiosensor can be designed in a short sequence to detect gene expression activities at a single-cell level. This ability enables us to correlate the Dll4 mRNA expression to the cell behaviors and to monitor the dynamic gene expression profiles during osteogenic differentiation.

Notch signaling pathway is an evolutionarily conserved intercellular pathway plays a substantial role in regulating stem cell behaviors in numerous tissues and organisms. [42, 43] This pathway plays a crucial role in lineage-specific differentiation of stem cells, and multiple stages of organismal development [44]. There are four Notch receptors (Notch1, Notch 2, Notch 3 and Notch 4) and five Notch ligands (Delta-like 1 (Dll1), Dll3, Dll4, Jag1, and Jag2). It has been reported that in recent years Notch signaling is involved in MSCs differentiation. For instance, Semenova *et al*. reported that Notch signaling regulates osteogenesis is dose-dependent, low-dose activation enhances osteogenic differentiation, while high-dose activation leads to apoptosis.[45] Cao *et al*. studied the involvement of Notch receptor Notch1 and Notch ligand Dll1 in osteogenic differentiation.[21] They observed that Notch1 inhibition reduced ALP activity during BMP-induced osteogenic differentiation of hMSCs *in vitro*. In contrast, it has been reported that inhibition of Notch signaling promotes adipogenic differentiation of MSCs,[46] indicating that Notch involvement is lineage-dependent during MSCs differentiation. Although numerous studies have demonstrated the involvement of Notch receptors or ligands, the involvement of Notch ligand Dll4 in osteogenic differentiation has not been explored. Here, for the first time, we studied the involvement of Notch ligand Dll4 during hMSCs osteogenic differentiation in 2D and 3D microenvironment. By exploiting an LNA/DNA nanobiosensor, we examined the dynamic expression of Dll4 mRNA during osteogenic differentiation. Our results showed Dll4 mRNA is highly expressed in differentiated hMSCs and has a dynamic change at the early stage of differentiation, **Fig. 3**. This is consistent with previously reported data that hMSCs experience different phases of cell-fate determination.[28] Our results showed that Dll4 mRNA reached the highest expression after 5 days of induction, indicating the involvement of Notch ligand Dll4 in early lineage progression. We further investigated the correlation of Dll4 mRNA expression and osteogenic differentiation by evaluating the early osteogenic differentiation marker, ALP activity. Our results indicate that Dll4 mRNA is a molecular signature of differentiated hMSCs. We next investigated Notch1-Dll4 signaling in regulating osteogenic differentiation in a 2D microenvironment. Disruption of Notch signaling using γ-secretase inhibitor DAPT reduced ALP activity during osteogenic differentiation and decreased the Dll4 mRNA expression. However, Jag1 treatments did not show a significant influence on osteogenic differentiation, which is consistent with previous results. It has been reported that Jag1 regulates osteogenic differentiation is dose-dependent, indicating the different effects of Jag1 treatments. [45] Moreover, the effects of Notch1-Dll4 signaling on hMSCs osteogenic differentiation in 3D spheroids were evaluated. Our results indicate that Notch1 inhibition enhanced the proliferation of hMSCs spheroids and reduced osteogenic differentiation with increased spheroid size and decreased ALP activity. Overall, our results suggested that hMSCs osteogenic differentiation is regulated by Notch1-Dll4 signaling, as supported by pharmacological treatments. Further investigation is required to clarify the involvement of other members of Notch family in the regulation of osteogenic differentiation.

## Conclusion

In this study, an LNA/DNA nanobiosensor is exploited to monitor the mRNA expression of hMSCs during osteogenic differentiation. Unlike conventional techniques, such as RT-PCR, which requires a large number of cells, this LNA/DNA biosensor detects gene expression at the single cell level. Another advantage of this LNA/DNA biosensor is that it doesn’t need cell lysis for fixation, permitting the investigation of dynamic gene expression during osteogenesis. Here, we investigated Notch1 ligand Dll4 mRNA expression dynamics during osteogenic differentiation. Our results showed that Dll4 expression levels correlate with the phenotypic and differentiation markers of hMSCs during osteogenesis. The ability to monitor Dll4 mRNA in living cells enables us to identify the regulatory role of Notch signaling. This study revealed that Notch1-Dll4 signaling is involved and active during osteogenic differentiation. Inhibition of Notch signaling by a γ-secretase enzymatic inhibitor attenuates the ALP enzymatic activity and decreases Dll4 mRNA expression. Our results also demonstrate that Dll4 mRNA levels correlated with early osteogenic markers (ALP activity), indicating that Dll4 mRNA is a molecular signature of differentiated hMSCs. In the absence of Notch inhibitor, hMSCs, cultured in osteogenic medium, differentiated towards the osteoblast phenotype, has high Dll4 expression. In conclusion, the results of this study add new information concerning osteogenic differentiation of hMSCs and the involvement of Notch1-Dll4 signaling. Further studies may elucidate the mechanisms underlying Notch1-Dll4 signaling in regulating osteogenesis.

## Materials and Methods

### Cell culture and reagents

Human mesenchymal stem cells (hMSCs) were acquired from Lonza and maintained in mesenchymal stem cell basal medium (MSCBMTM) with GA-1000, L-glutamine, and mesenchymal cell growth supplements. The cells were cultured in a humidified incubator at 37 °C with 5% CO_2_ and passaged using 0.25 Trypsin-EDTA (Invitrogen). The cell culture medium was replaced every three days. hMSCs from passages 3-7 were used in the experiments. DAPT was purchased from Sigma Aldrich (DAPT ≥98% (HPLC), solid, D5942). To induce osteogenic differentiation, hMSCs were seeded at a density of 1× 10^4^ cells/mL with a volume of 500 μL in 12 well-plates. The basal medium was replaced with osteogenic induction medium after two days of cell seeding. Osteogenic induction medium were changed every two days. For studying Notch1-Dll4 signaling, hMSCs were treated with 20 μM γ-secretase inhibitor DAPT or 20 μM Jag1 peptide (AnaSpec, 188-204) after osteogenic induction. It is noted that both DAPT and Jag1 were added daily. Images were taken after 3 days, 5 days, 7 days, and 10 days of induction, respectively.

### 3D spheroids formation and culture

We fabricated 3D self-organized spheroids of hMSCs using hanging drop approach [41]. A process flow illustrating the 3D hMSCs spheroids formation is shown in **Fig. S5**. First, hMSCs were harvested for spheroid formation using 0.25% trypsin-EDTA. The cells were then centrifuged for 5 minutes at 200 G to form a cell pellet. Then trypsin were aspirated and replaced with fresh medium with 20% methylcellulose (Sigma Aldrich, M0512) added. The final concentration of the cell solution is 4 × 10^5^ cells/mL. Then 20 μL drops of the mixture were placed on the underside of a petri dish lid, with 5 mL of 1x PBS inside the dish to prevent droplet evaporation. In our experiments, 75 droplets were seeded per lid. Spheroids were then incubated as hanging drops in the 37 °C, 5% CO_2_ incubator for 72 hours. Next, the spheroids were harvested using open-end 1 mL tips to wash off the spheroids from the lid at a small amount of basal growth medium and keep the lid inclined to ensure the spheroids slipped over. Then all the spheroids were transferred into a pre-sterilized 1.5 mL microcentrifuge tube (Labnique, Hunt Valley, MD). As the spheroids are gathered in the bottom of the centrifuge tube, aspirate the medium as much as possible, then mix spheroids in growth factor reduced Matrigel (Corning, 356234). The 3D spheroids mixed in Matrigel were then placed in 24-well plates. For Dll4 mRNA detection, hMSCs were transfected with LNA/DNA probe before forming spheroids. For evaluating ALP activity in 3D spheroids, ALP staining was performed after 10 days of osteogenic induction.

### LNA Probe Design

An LNA probe is a 20-base pair nucleotide sequence with alternating LNA/DNA monomers. A fluorophore (6-FAM) was labeled at the 5’ end for fluorescence detection. The design process of LNA probes was reported previously [47-50]. Briefly, the LNA probe was designed to be complementary to the target mRNA sequence with a 100% match. The binding affinity and specificity were optimized using mFold server and NCBI Basic Local Alignment Search Tool (BLAST) database. A quencher is a 10-base pair nucleotide sequence with LNA/DNA monomers that is complementary to the 5’ end of the LNA probe. An Iowa Black RQ was labeled at the 3’ end of the quencher probe. The Dll4 probe was designed based on target mRNA sequences, **Supplementary Tab. S1**). A random probe, lacking a complementary mRNA, was designed as the negative control. The LNA probe was synthesized by Exiqon Inc. The quencher sequence and corresponding target DNA sequences were synthesized by Integrated DNA Technologies Inc. (IDT).

### Double-stranded LNA probe preparation

To prepare the LNA/DNA nanobiosensor, the LNA probe and quencher were initially prepared in 1x Tris EDTA buffer at a concentration of 100 nM. The LNA probe and quencher were mixed and incubated at 95 °C for 5 minutes and cooled down to room temperature over the course of 2 hours. To optimize quenching efficiency, the fluorescent probe and quencher probe were prepared in a number of different ratios to obtain high signal-to-noise ratio. The quenching efficiency was evaluated by measuring fluorescence intensity using a fluorescence microplate reader (BioTek, Synergy 2). The optimized LNA probe and quencher mixer can be stored in a refrigerator for up to 7 days. The prepared double-stranded LNA/DNA probe (100 nM) was transfected into hMSCs using Lipofectamine 2000 in opti-MEM from Invitrogen to determine the target gene expression. Gene expression was evaluated by measuring the fluorescence intensity of cells transfected with LNA/DNA probes.

### Alkaline Phosphatase Activity (ALP) Staining

To quantify hMSCs osteogenic differentiation, cells were stained for alkaline phosphatase (ALP) using the alkaline phosphatase kit (Sigma-Aldrich) using a modified protocol. First, the staining solution were prepared by mixing fast red violet solution, naphthol AS-BI phosphate solution, and water at a ratio of 2:1:1. Next, hMSCs were fixed using 4 % cold Paraformaldehyde (PFA) for 2 minutes which enabled the maintenance of the ALP activation. The staining solution was then added to fixed cells for 15 minutes under room temperature and protected from light. For nucleus staining, Hoechst 33342 staining solution was prepared in 1x PBS at 1:2000 dilution and added to cells for 15 minutes. The cells were then washed three times with 1x PBS, 15 minutes each time, before taking images.

### Imaging and Statistical Analysis

Images were captured using ZOE Fluorescent Cell Imager (BIO-RAD) or Nikon TE 2000. All fluorescence images were taken with the same setting for comparison. Data collection and imaging analysis were performed using NIH ImageJ software. Experiments were repeated at least three times, and over 100 cells were quantified for each group. For 3D spheroids, at least 20 spheroids were analyzed for each group. Results were analyzed using independent, two-tailed Student *t*-test in Excel (Microsoft). *P* < 0.05 was considered statistically significant.

## Data availability

The data that support the findings of this study are available from the corresponding author on reasonable request.

## ACKNOWLEDGMENT

This work is supported by NSF CAREER Award (Award number: 2143151) and NASA CT Space Grant Consortium Faculty Research Grant (Award Number: P-1558). Y Zhao is supported by the Provost Graduate Fellowship.

## Author contributions

Y.Z and S.W conceived the initial idea of the study. Y.Z, R.Y, Z.B, and K.R performed the experiments. Y.Z and S.W contributed to the experimental design and data analysis. Y.Z and S.W wrote the manuscript with feedback from all authors.

## Additional information

Competing financial interests: The authors declare no competing financial interests. Supplementary information: Fig. S1 – Fig. S5, Tab. S1.

## Data availability statement

All data that support all the experimental findings in this article are available from the corresponding author on reasonable request from individuals affiliated with research institutions for research use only.

