## Supplementary Materials for "Probing Notch1-Dll4 Signaling in Regulating Osteogenic Differentiation of Human Mesenchymal Stem Cells using Single Cell Nanobiosensor"

**Fig. S1.** Random probe expression tracking of hMSCs during osteogenesis for 7 days **.**

**Fig. S2.** Dll4 mRNA expression tracking of hMSCs during osteogenesis for 7 days..

**Fig. S3.** Representative bright field and fluorescence images of hMSCs after 5 days of osteogenic induction.

**Fig. S4.** Representative bright field and fluorescence images of hMSCs after 5 days of osteogenic differentiation under different treatments.

**Fig. S5.** Schematic illustration of hMSCs 3D spheroid formation.

**Tab. S1.** LNA/DNA probes and quencher sequences

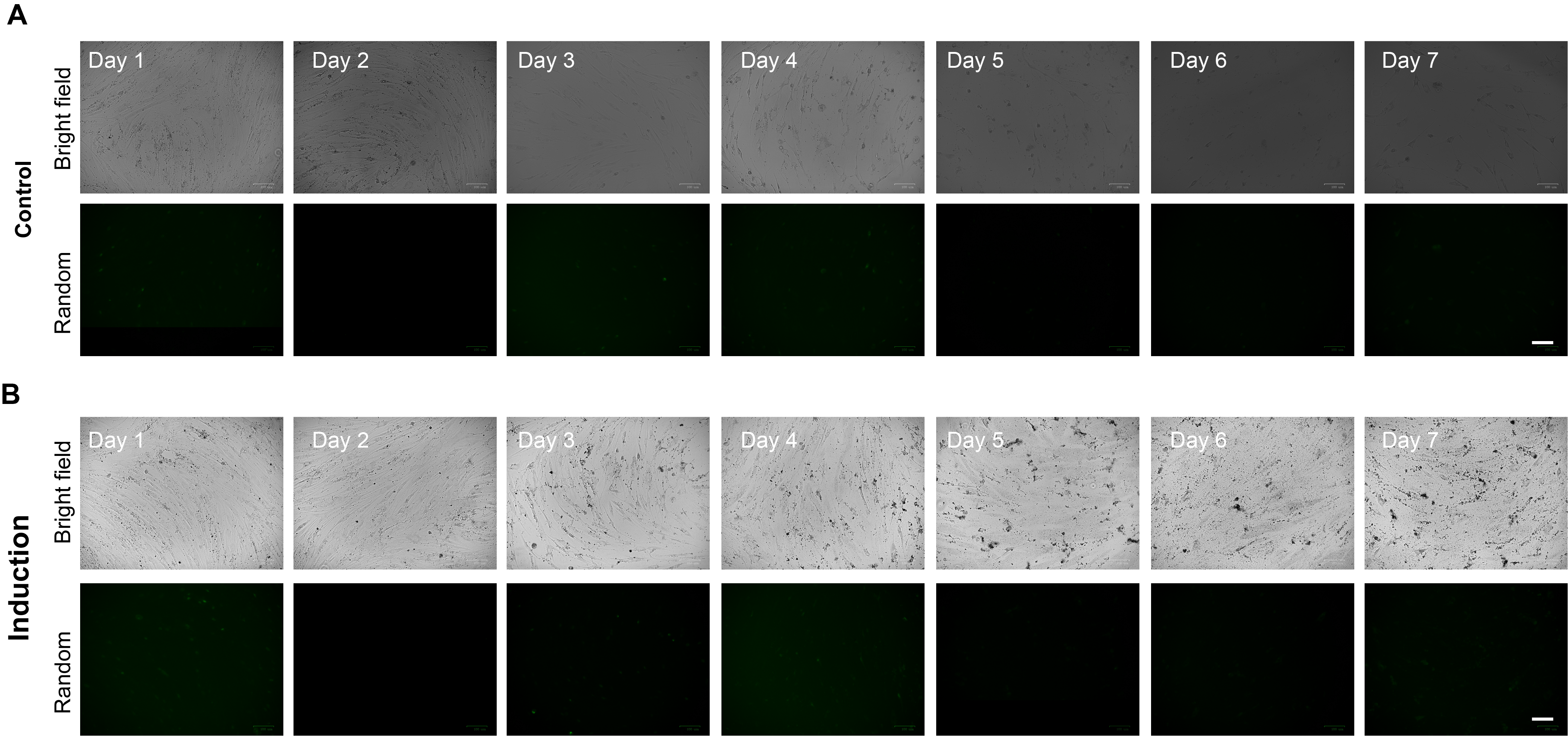

**Fig. S1**. Random probe expression tracking of hMSCs during osteogenesis for 7 days. hMSCs were cultured in basal medium **(A)** and osteogenic induction medium **(B)** for 7 days, respectively. Green: random probe. Scale bar: 100 μm.

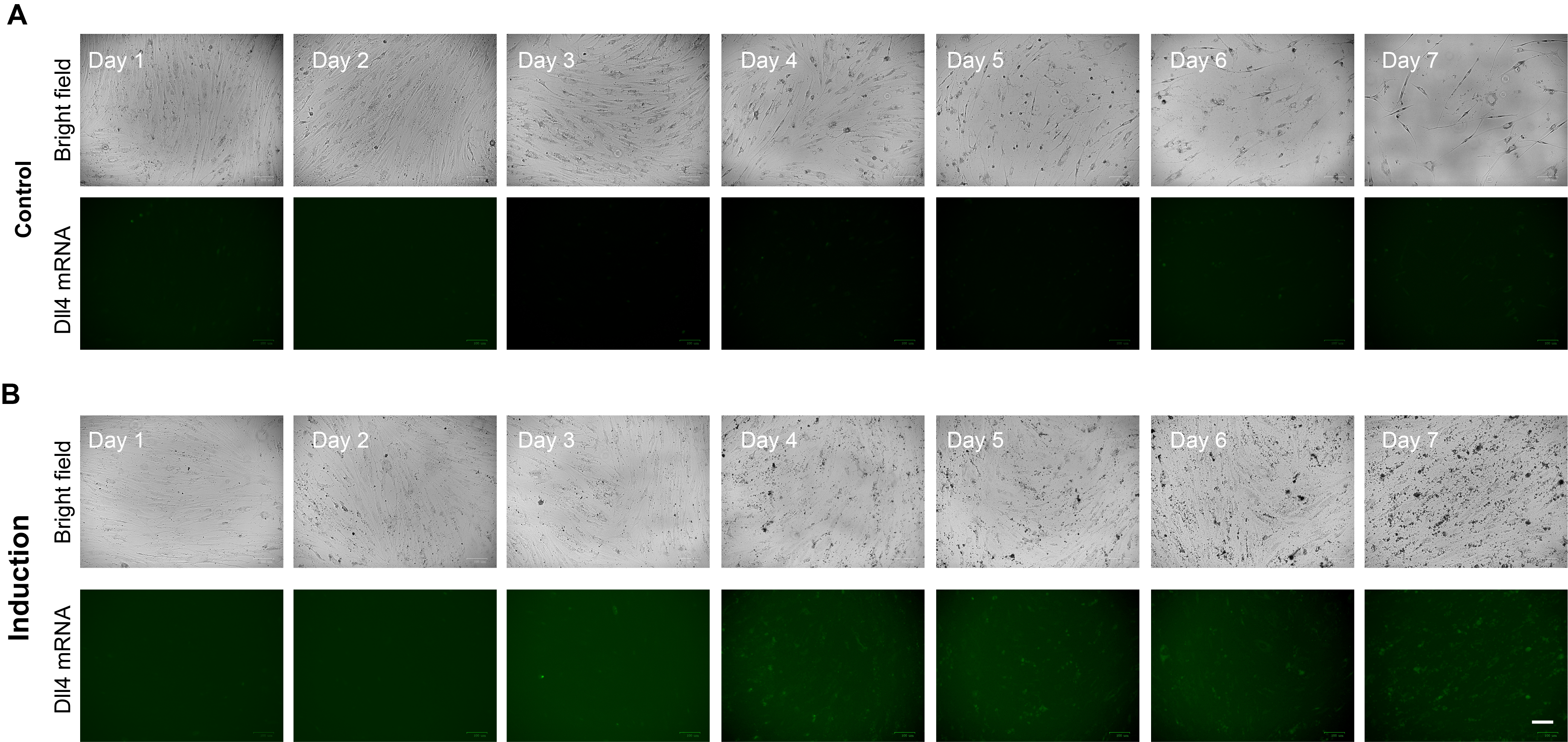

**Fig. S2.** Dll4 mRNA expression tracking of hMSCs during osteogenesis for 7 days. **(A)** Representative bright field and fluorescence images of hMSCS cultured in basal medium. **(B)** Representative bright field and fluorescence images of hMSCS cultured in osteogenic induction medium. Green: Dll4 mRNA. Scale bar: 100 μm.

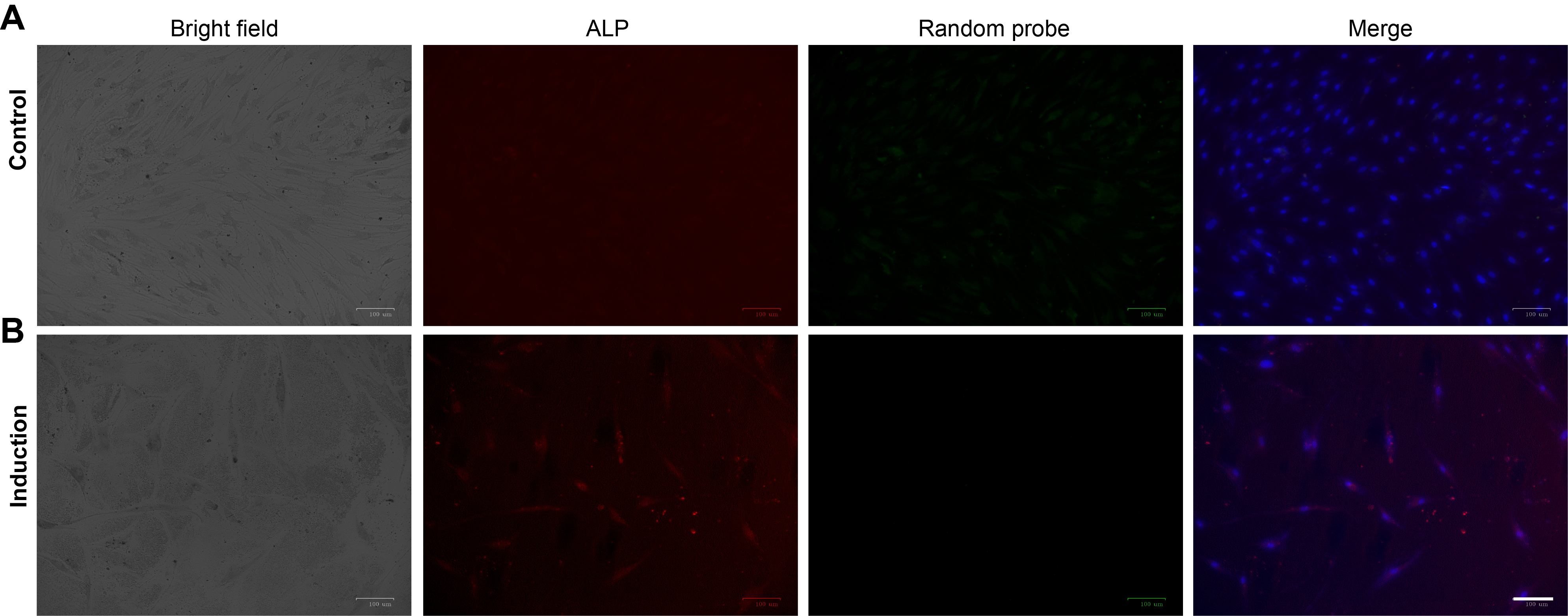

**Fig. S3**. Representative bright field and fluorescence images of hMSCs after 5 days of osteogenic induction. hMSCs were transfected with a random probe and stained with ALP and Hoechst 33342. **(A)** Representative images of hMSCs cultured in basal medium. **(B)** Representative images of hMSCs cultured in osteogenic induction medium. Green: random probe; red: ALP; blue: Nucleus. Scale bar: 100 μm

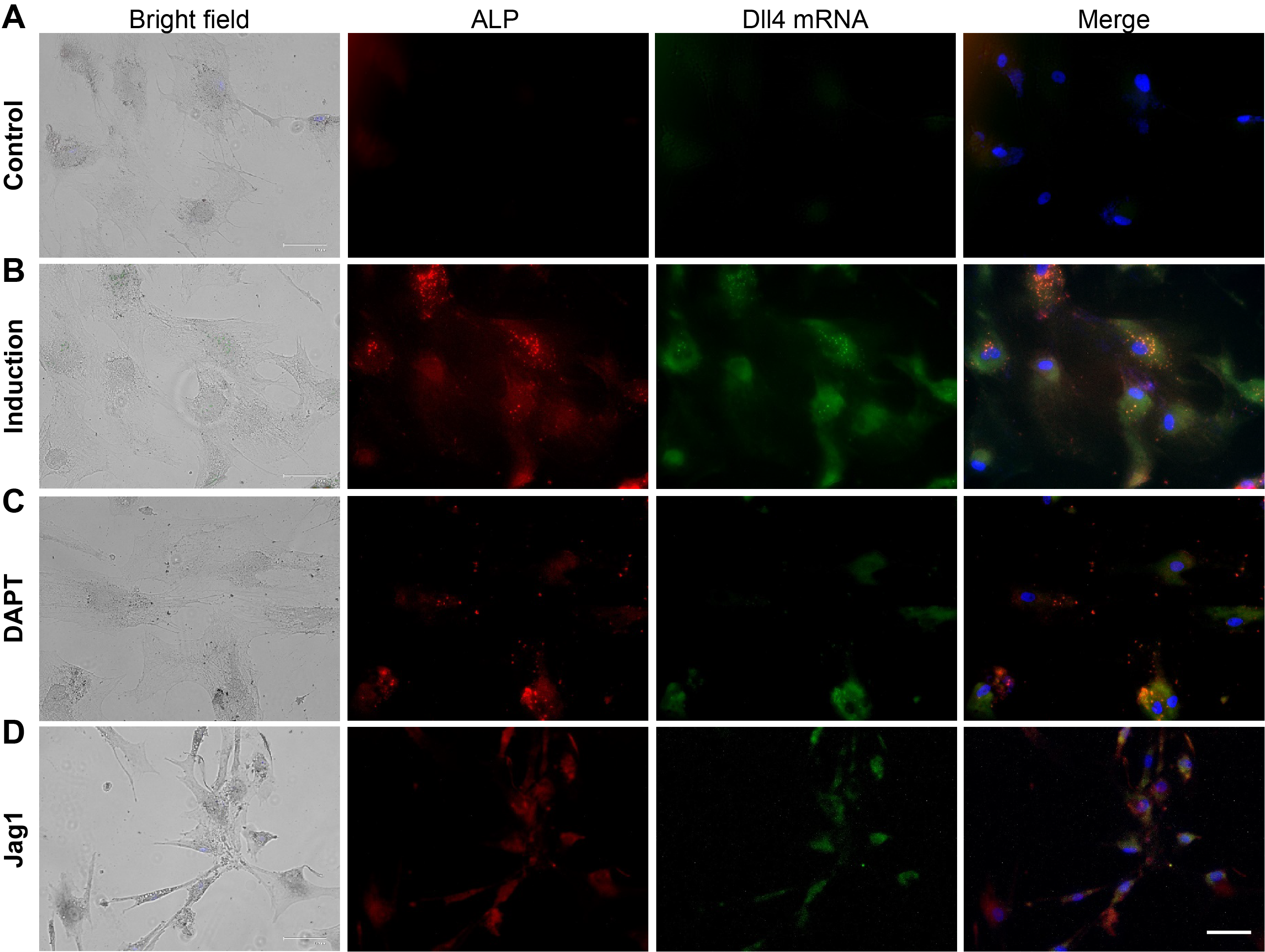

**Fig. S4**. Representative bright field and fluorescence images of hMSCs after 5 days of osteogenic differentiation under different treatments. **(A)** Control group; hMSCs were cultured in the basal medium without treatments. **(B)** Induction group; hMSCs were cultured in osteogenic induction medium without treatments. **(C)** DAPT group, hMSCs were cultured in induction medium and treated with DAPT at a concentration of 20 μM for 5 days. **(D)** Jag1 group, hMSCs were cultured in induction medium and treated with Jag1 peptide at the concentration of 40 μM for 5 days. Green: Dll4 mRNA; red: ALP; blue: Nucleus. Scale bar: 100 μm.

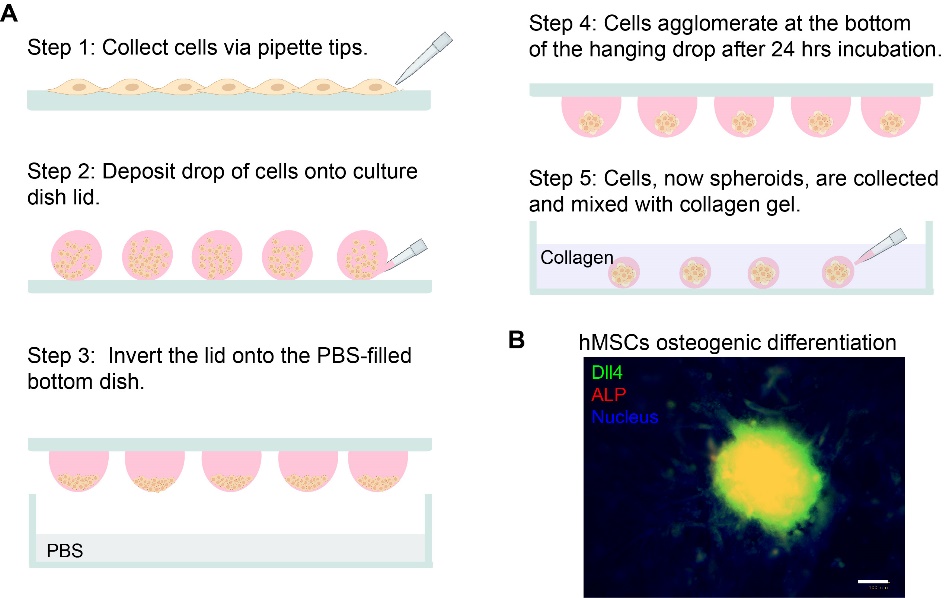

**Fig. S5**. Schematic illustration of hMSCs 3D spheroid formation. **(A)** Process flow of spheroid formation using a hanging drop method. Cells were harvested and seeded at a concentration of 8x10^3^ cells/drop on a culture dish lid. Spheroids were then incubated as hanging drops for 72 hrs, and then transferred into the Matrigel. **(B)** Representative images of 3D spheroids after 10 days of osteogenic differentiation. Green: Dll4 mRNA; red: ALP; blue: Nucleus. Scale bar: 100 μm.

**Tab. S1.** LNA/DNA probes and quencher sequences

| Name | | Sequence (5’-3’) | Fluorophore |
| --- | --- | --- | --- |
| Dll4 mRNA | Donor | +AA +GG +GC +AG +TT +GG +AG +AG +GG +TT | /56-FAM |
|  | Quencher | +TT +CC +CG +TC +AA | /3-Iowa BlackFQ |
|  | Target | AA CC CT CT CC AA CT GC CC TT |  |
| Random | Donor | +AC+GC+GA+CA+AG+CG+CA+CC+GA+TA | /56-FAM |
|  | Quencher | +TG +CG +CT +GT +TC | /3-Iowa BlackFQ |
|  | Target | TA TC GG TG CG CT TG TC GC GT |  |

* + represents LNA monomer
